# Existence of blood circulating immune-cell clusters (CICs) comprising antigen-presenting cells and B cells

**DOI:** 10.1101/2022.01.26.477834

**Authors:** Sangwook Bae, Yonghee Lee, MyeongHo Kang, Jinsung Noh, Dongyup Seo, Hanna Seo, Sumin Lee, Sunghoon Kwon

## Abstract

Cell-to-cell physical interactions are involved in almost every physiological processes in multicellular organisms. Although the dynamics of these interactions could be highly diverse and complex in many circumstances, certain cell-to-cell interactions among immune cells have been well studied due importance in understanding disease pathogenesis and immune therapy development^1^. Dendritic cells (DCs) and B cells are directly involved in adaptive immune response against pathogens. Interaction mechanism between these two celltypes is well-known to occur in germinal centers either indirectly via helper T (Th) cells or directly via cell contact. However, there are animal *in vitro* and *in vivo* evidence that such direct DC-to-B cell contact can occur outside germinal centers like in peripheral blood or collagen matrix and display antiviral immune-related activity^2,3^. Here, we provide evidence that certain types of antigen presenting cells (APCs) can form robust cell clusters with B cells and circulate in blood. From healthy human blood immune single cell RNA-seq datasets, we detected APC subpopulations (0.34±0.19% of total PBMCs) that were also enriched with well-known naïve B cell markers. We visually observed DC:B doublets and multiplets (∼0.69% of total live PBMCs) in wildtype mouse blood using flow cytometry and microscopic imaging, thus proving the existence of circulating immune-cell clusters (CICs) composed of APCs and B cells. BCR repertoire of these healthy mouse CICs were similar to circulating B cells. Noticeably, frequency of these APC:B CICs were higher COVID-19 patients than healthy donors and their B cell subtype composition (e.g. naïve, plasmablast, IgM^+^, IgG^+^) varied with disease severity.

## Introduction

Recent progress in single cell sequencing technologies, spatiotemporal tissue analysis and other biochemical assays are now allowing researchers to investigate the complex cell-cell interaction networks that shape and control the multicellular organs and organisms. Many cell-cell communications are contact-independent mechanism (ligand secretion) but certain key physiological processes (neural synapse, tissue organization, innate and adaptive immune reaction etc.) require direct cell-cell interaction^4^. Understanding such direct cell-cell interactions are becoming of great importance for developing state of the art therapeutics such as immune checkpoint inhibitors^5^ and CAR-T therapy^6^. Analyzing such direct cell-cell interactions can be done with a number of biochemical strategies (e.g. cellular imaging and proximity tagging)^1^ but the most recently prominent ones are based on single cell sequencing and spatial sequencing^7^. Such direct cell-cell interaction assays can also act as validation platform for intercellular communication network studies that are based on ligand-receptor network analysis. They also enable exploratory analyses, like those demonstrated with recently developed platforms like PIC-seq^8^ and CIM-se^9^, of previously unrecognized interactions among subtypes of immune cells and solid tissue cells.

Dendritic cell (DC), as the most effective antigen presenting cell (APC), is well known celltype to interact with T cells to enable adaptive immune against the presented antigen. They regulate B cell-based humoral immune usually by indirect mechanisms where Th cells as used as mediators. But there are also known cases of direct DC-to-B cell contact happening in lymphoid organs^10-12^. For example, previous studies showed that myeloid-derived DCs can regulate B-cell differentiation via cell-to-cell contact^13^. Another study demonstrated that autoantigen-stimulated B cells can directly interact with plasmacytoid dendritic cells (pDCs) and upregulate their INFα production^11^. Also, these IFNα-producing pDCs stimulated differentiation and immunoglobulin production of the contacting B cells, in the absence of T cell help^10, 14^. Another study showed by direct cell-cell contact pDCs can stimulate autoreactive B cell activation, autoantibody production, and survival *in vivo* through TLR and BCR stimulation^12^. This implies potential etiopathogenic involvement of such pDC-B interaction in prolonged type 1 IFN activation often present among Systemic lupus erythematosus (SLE) or other autoimmune disease patients^11^. Overall, these studies focused on *in vitro* co-cultures and lymphoidal organs. However, there is little evidence about whether cell clusters comprising DCs and B cells exist in peripheral blood.

Here, we present evidence of the existence of blood-circulating immune cell clusters (CICs) comprising APCs and B cells cell. From publically available scRNA-seq results of human peripheral blood mononuclear cells (PBMC), we repeatedly detected data points composed of a mixture of B cells with monocytes or dendritic cells. Such APC:B CICs were observable in PBMCs of human and mouse. We confirmed the existence of DC:B CICs in blood samples from wildtype mice using flow cytometry and microscopy. These DC:B CICs contained BCR repertoire similar to blood circulating B cells which implies that this DC-to-B cell interaction occurs in a more general, BCR-independent manner. These B cell marker-enriched APCs were mostly CD14+ Monocytes or CD1c+ DCs and rarely detected in other APC types. This is unlikely to have happened if they were artifacts of cell free RNA contamination. They were detected in scRNA-seq data of flow-cytometry sorted single APCs and EDTA-treated single APCs. So it is also unlikely that they have occurred from accidental mixture of two independent cells or nonspecifically charge-bound cell doublets. Although whether Th cells are involved in the initial formation of these APC:B CICs is uncertain, we concluded that, based on our observations, there are CICs composed of directly contacting APCs and B cells circulating in blood.

## Materials and Methods

### Antibodies

For the DC-B, B cell FACS and BCR repertoire analysis, anti-Mo CD3-FITC (17A2, BioLegend), anti-Mo CD49b-PE (DX5, BioLegend), anti-Mo CD11c-PE-Cy7 (N41B, eBioscience), anti-Mo CD19-APC (1D3, BioLegend) and anti-Mo I-A/I-E-APC (M5/114.15.2, BioLegend) was used. For the DC-B FACS and microscope imaging experiment, anti-Mo CD3-FITC (17A2, BioLegend), anti-Mo NK1.1-APC-Cy7 (PK 136, BioLegend), anti-Mo I-A/I-E-APC (M5/114.15.2, BioLegend), anti-Mo CD11c-Pacific Blue (N418, BioLegend), anti-Mo CD19 biotin antibody (MagniSort, Invitrogen) and Streptavidin-PE (S866, Thermofisher) was used.

### Mouse cell isolation and FACS

Wild-type C57BL/6 were maintained in specific pathogen-free (SPF) animal care facilities at Sungkyunkwan University (SKKU) following the Institute/University Animal Care and Use Guidelines. Mouse PBMCs were isolated from the blood of SPF mice using Ficoll (GE Healthcare, Little Chalfont, UK) density gradient centrifugation and PBS washing. Mouse lymphoidal cells were isolated from inguinal lymph nodes (iLNs) by cell straining and PBS washing. For immunofluorescence staining, cells were stained in FACS buffer at 4°C for 20 min with the appropriate antibodies. After washing with FACS buffer, cells were analyzed by FACSCantoII (BD Biosciences) and FACS DIVA software. Compensations were performed with single-stained UltraComp eBeads (Affymetrix) or cells. DC-B pair gating was done as follows: FVD-(live cells), FSC-W^high^ (Doublet), lineage (CD49b or NK cell)-negative, MHC-II+CD11c+ (DC) and CD19+ (B cell).

### Fluorescence microscopy

Microscope imaging of DC-B CICs were done using Nikon Ti-E with Pan Fluor 20x (NA 0.45) Ph1 objective lens, Hamamatsu digital camera. Standard AT series filter cubes were used for filtering signals from anti-Mo CD19-biotin-streptavidin-PE and anti-Mo CD11c-Pacific Blue.

### BCR RT-PCR and NGS library preparation

RNA from FACS sorted DC-B cells or B cells was purified using Qiagen RNeasy mini kit. For reverse transcription, purified RNA input of 40 to 100ng was used per sample. Purified RNA was mixed with reverse transcription primer (final 5μM) and dNTP (final 0.77mM) and denatured at 72°C for 3min and placed on ice for 2 min. 5x buffer, DTT, RNAseOUT, and Superscript IV reverse transcriptase was added as described in the manufacturer’s guide. Reverse transcription was done at 55°C for 10 minutes and heat inactivated (80°C 10min). First strand cDNA was purified using 1.8x AMPure XP bead purification as described in the manufacturer’s guide and eluted in nuclease free water. Second strand cDNA was synthesized using KAPA HiFi polymerase single cycle PCR with 4μM second strand synthesis primer mix. PCR was done by 95°C 3min, 98°C 30sec, 60°C 30sec, and 72°C 5.5min. Second strand cDNA was purified with 1x AMPure XP bead purification and eluted in nuclease free water. Final NGS library PCR was done using KAPA HiFi polymerase and 0.4μM Nextera XT index primers. PCR protocol was as follows: 5°C 3min; 8 cycles of 98°C 30sec, 62°C 30sec, and 72°C 30sec; 72°C 5min. Library was purified with 1x AMPure XP bead and eluted in nuclease free water. Library was sent to SNUH for QC and was sequenced using NovaSeq (illumina).

### BCR NGS data analysis

The in-house preprocessing on NGS raw reads was conducted as described previously^15^, which was composed of four steps; paired-end merge, quality filtering, error correction, and region annotation. First, all raw paired FASTQ files were subjected to paired-end merge process using the PEAR software ^16^ with default parameters. Then, quality filtering on merged files were done with the Q20P95 option, which extracted the reads if more than 95% of bases had a Phred quality score of more than 20. For the error correction, UMI-based error-correction, which could exclude errors by extracting consensus sequence from the each cluster of reads having the same UMI was applied. In region annotation process, The V/J gene and complementary determining regions (CDRs; CDR1, CDR2, and CDR3) were annotated using the IgBLAST tool (version 1.17.1)^17^, with the Ig germline database of the C57BL/6 mice (Mus musculus) acquired from the IMGT database^18^. Reads which were not functional, such as having stop codon or out-of-frame, were excluded. From functional reads which passed whole preprocessing steps, three features of BCR repertoire were computed and compared; the abundance of isotypes, clonality, and the number of somatic hypermutations (SHMs) in the V gene. The isotype of each sequence was computed as the germline C gene of highest similarity using the BLAST tool^19^. The abundance of each isotype was defined as the number of sequences for each isotype divided by the total number of sequences. To measure clonality, which is the measure of unevenness in the distribution of BCR sequences, the Gini index was calculated using the frequencies of sequences. The number of SHM was computed by counting the mutations in the sequence based on its related germline V gene. The whole comparative figures were drawn using ggplot2 package in R (version 3.6.3), and the Wilcoxon rank sum test was applied to test the significance of the difference in the number of SHM between APC-interacting and non-interacting B cells.

### scRNA-seq data analysis

Seurat (v 3.1)^20^ was used for integrating multiple scRNA-seq datasets and downstream analyses. Dataset integration algorithm was based on canonical correlation analysis (CCA) and matched mutual neighboring (MNN). The CCA and L2-normalization was effective for removing batch effects while MNN was used for dimensionality reduction. MNN searches for cell pairs (anchors) that encode the the cellular relationships across datasets that serves as basis for all subsequent integration analyses. PIC-seq analysis was done with algorithms provided by the original pape^8^. For this, DC geneset and B cell geneset was extracted from DC clusters and B cell clusters present in 10x Genomics healthy PBMC scRNA-seq data using FindMarkers() function. Hierarchical clustering was done with hclust() function with “ward.D2” method. Publically available scRNA-seq datasets from Gene Expression Omnibus (GEO) used for this study are as follows: GSE169346, GSE150728, GSE94820. Healthy donor PBMC scRNA-seq datasets (10x Genomics) was also used (https://www.10xgenomics.com/resources/datasets).

## Results

### Observation of blood APCs containing B cell signatures in human PBMC scRNA-seq data

From the publically available 10x Genomics’ healthy human PBMC scRNA-seq dataset, we detected a small cluster of APCs that also displayed B cell signatures (**Figure 1.A**). For this analysis, we selectively removed lymphocyte clusters which were dominated by either one of CD3E^+^ T cells, CD19^+^ B cells or CD56^+^ (NCAM1^+^) NK cells and had no significant mixed signatures of myeloid cells. We then analyzed the remaining myeloid cells including DCs (HLA-DR^+^CD11c^+^CD16^−^), monocyte (HLA-DR^+^CD14^+^CD16^-^) and monocyte-derived DC (mo-DC) (HLA-DR^+^CD11c^+^CD14^+^CD226^+^) and macrophages (HLA-DR^+^CD11c^+^CD1c^−^CD16^+^) according to published studies^21-23^. Among DC subtypes, clusters of CD1c^+^ DCs and IL3RA^+^CD226^+^ pDC were present in these datasets while CLEC9A^+^ DC were hardly detectable. CD141^+^ DCs existed but the cluster was ambiguous due to overall low level of CD141 detected in these datasets. Noticeably, we detected a unique cluster that contained cells with high expression of B cell markers (e.g. CD19 CD20 or MS4A1) as well as monocyte or DC markers (CD11c) (**Figure 1.B**). This mixed-phenotype cluster contained cells with high levels of CD14 and low expression of CD16. We designated this cluster as APC:B.

**Figure 1.**
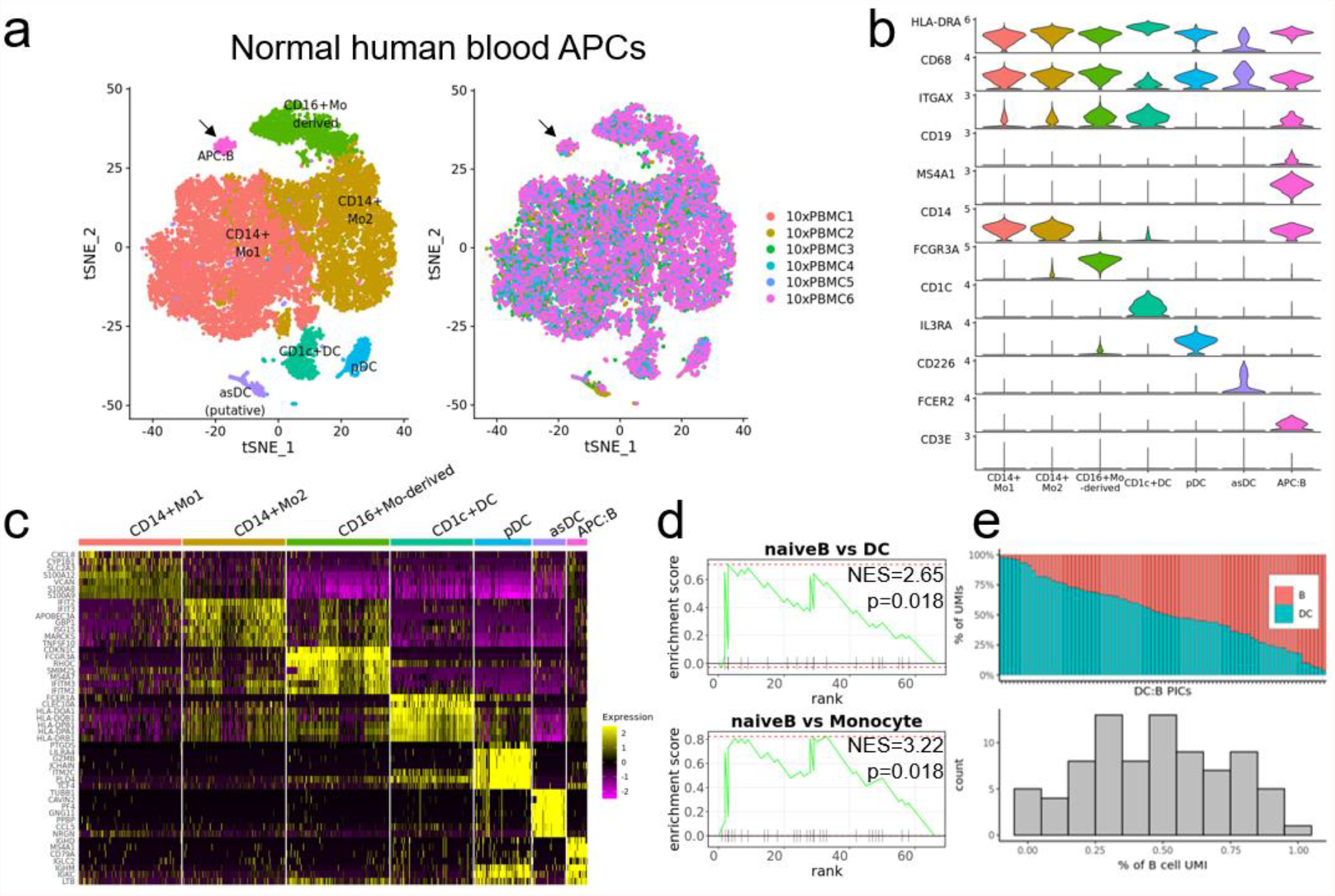
Existence of human blood circulating APCs containing B cell signatures. (a) t-SNE plot of healthy human blood APCs. (b) Violin plot of APC and B cell marker gene expression among different celltypes. Other than CD19 and MS4A1 are APC celltype-specific gene. (c) Heatmap of differentially expressed genes of different APC clusters. (d) GSEA analysis of the APC:B cluster-specifically enriched genes against naïve B enriched gene set. Upper panel is result against gene panel of genes upregulated in B cells compared to DCs. Lower panel is result against gene panel of genes upregulated in B cells compared to monocytes. (e) PIC-seq analysis result of APC:B cluster. Upper panel shows transcript composition of each data point predicted to have come from either DC or B cells. Lower panel is a histogram of percentage of B cell-derived transcripts from each data point. DC-enriched genes and B cell-enriched genes used for this analysis were extracted from the B cell clusters and DC clusters in the 10x Genomics healthy PBMC scRNA-seq dataset.

One could argue that these B cell-specific transcripts in APC:B cluster are simple contamination of cell-free RNA from B cell debris. This is not likely because, as shown in violin plots (**Figure 1.B**), these B cell-specific transcripts were present dominantly in the APC:B cluster rather than being distributed evenly across different cell clusters or different cell types. It is highly unlikely that cell-free RNAs from B cells could explicitly mix into a certain celltype (i.e. CD14+ APC) at high concentrations during the droplet-based single cell sorting procedure. One could also argue these are independent APCs and B cells accidentaly encapsulated into the same droplet during droplet cell sorting. However, we detected APC:B-like signatures in scRNA-seq dataset generated from flow cytometry-sorted single human blood APCs (**Figure 2.B**). Appearance of this APC:B cluster was not a stochastic event because it was repeatedly observable among the 6 datasets with an average frequency of 0.34±0.19% of total PBMCs. Therefore, we hypothesized that this APC:B cluster is a cluster of blood CICs rather than being a spurious cluster formed by contamination of B cell’s cell-free mRNA or accidental cell mixture artefact.

**Figure 2.**
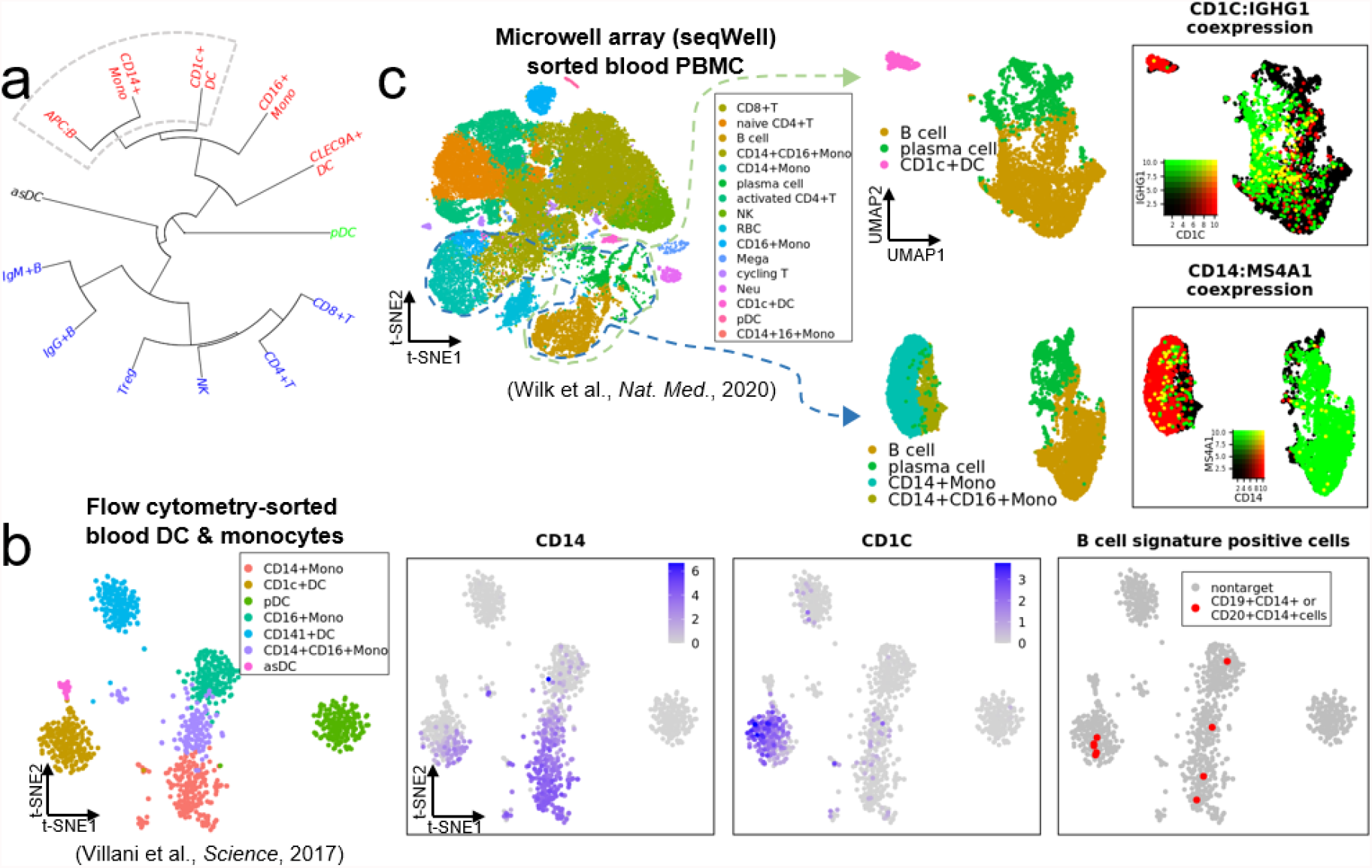
Existence of APC:B doublet signatures among flow cytometry-sorted or microwell array-sorted immune cells from healthy human blood. (a) Hierarchical clustering result of immune celltypes in healthy human PBMC data in figure 1. (b) t-SNE celltype and gene expression plots of flow cytometry-sorted healthy human blood DCs and monocyte single cell transcriptomes. Cells coexpressing B cell markers (CD19 or CD20) and CD14+ monocyte marker (CD14) was highlighted in red dots. (c) t-SNE celltype plot and two gene co-expression UMAP plots of microwell array-sorted healthy blood PBMCs. Gene expression UMAPs are selection of B cell and CD1c+ cells (upper panel) or B cell and CD14+ monocytes (lower panel). Cells co-expressing APC marker (red, CD1c for upper panel, CD14 for lower panel) and B cell marker (green, IGHG1 for upper panel, MS4A1 for lower panel) are highlighted in yellow.

### Gene expression characteristics of APC:B CICs

Gene set enrichment analysis (GSEA) showed that this APC:B cluster is enriched with B cell-specific gene expression patterns (**Figure 1.D**). And PIC-seq analysis showed that these DC:B CICs are indeed comprising both APC and B cell transcriptome (**Figure 1.E**). Hierarchical clustering showed that APC:B cluster has highest transcriptome similarity with CD14+ Monocytes, followed by CD1c+ DCs (**Figure 2. A**).

We then questioned whether these APC-binding B cells were of specific B cell subtype. Among the two B cell marker (IgM)-positive myeloid cell clusters - pDC cluster and APC:B cluster-the APC:B cluster showed high expression of IgM, IgD, and CD27 compared to the pDC cluster (**Figure S1. A, C, D**). This has similarity to a previous report, where it was shown that there are two types of circulating human B cells; marginal B cell-like IgM^high^IgD^+^CD27^+^ B cells and IgM^low^ naïve/memory B cells ^30^. When based on this report, it seems that the marginal B cell-like IgM^high^IgD^+^CD27^+^ B cells are enriched in the APC:B cluster, while IgM^low^ naïve/memory B cells are enriched in the pDC cluster. However, the pDC cluster cells were mostly negative for MS4A1 (CD20) (**Figure S1. D**), so whether any of these pDCs are bound to memory B cells is uncertain.

We observed that AIM2 was enriched in APC:B CICs compared to other DCs (**Figure S1. C**). AIM2 is a known mature B cell-specific marker and memory B cell-specific marker. The fact that AIM2 is enriched in APC:B cluster, other than B cell clusters, is in accordance with previous observations where naïve B cells differentiate into active or memory B cells on account of DC binding^10, 13, 14^. Although the small CLEC9A+ DC cluster was also positive for AIM2 expression, it is unlikely that the APCs in the APC:B cluster are CLEC9A+DCs because hierarchical clustering showed that the APC:B cluster was closest to CD14+ monocytes or CD1c+ DCs (**Figure S1. B, Figure 2.A**). But unlike most previous B cell-binding DC studies, B-cell binding DCs in our datasets were rarely derived from pDC clusters (**Figure S1 C, Figure 3. C**). This discrepancy could probably be explained by the lack of IFNα expression in these blood-circulating pDCs (**Figure S1. C**) which was shown to be necessary for differentiation and stimulation of the pDC-contacting B cells^10,14^. Future *in vitro* B cell co-culture with TLR9 ligand (e.g. CpG) PBMC-derived DC subtypes and TLR9 ligand (e.g. CpG) stimulation will address this hypothesis.

**Figure 3.**
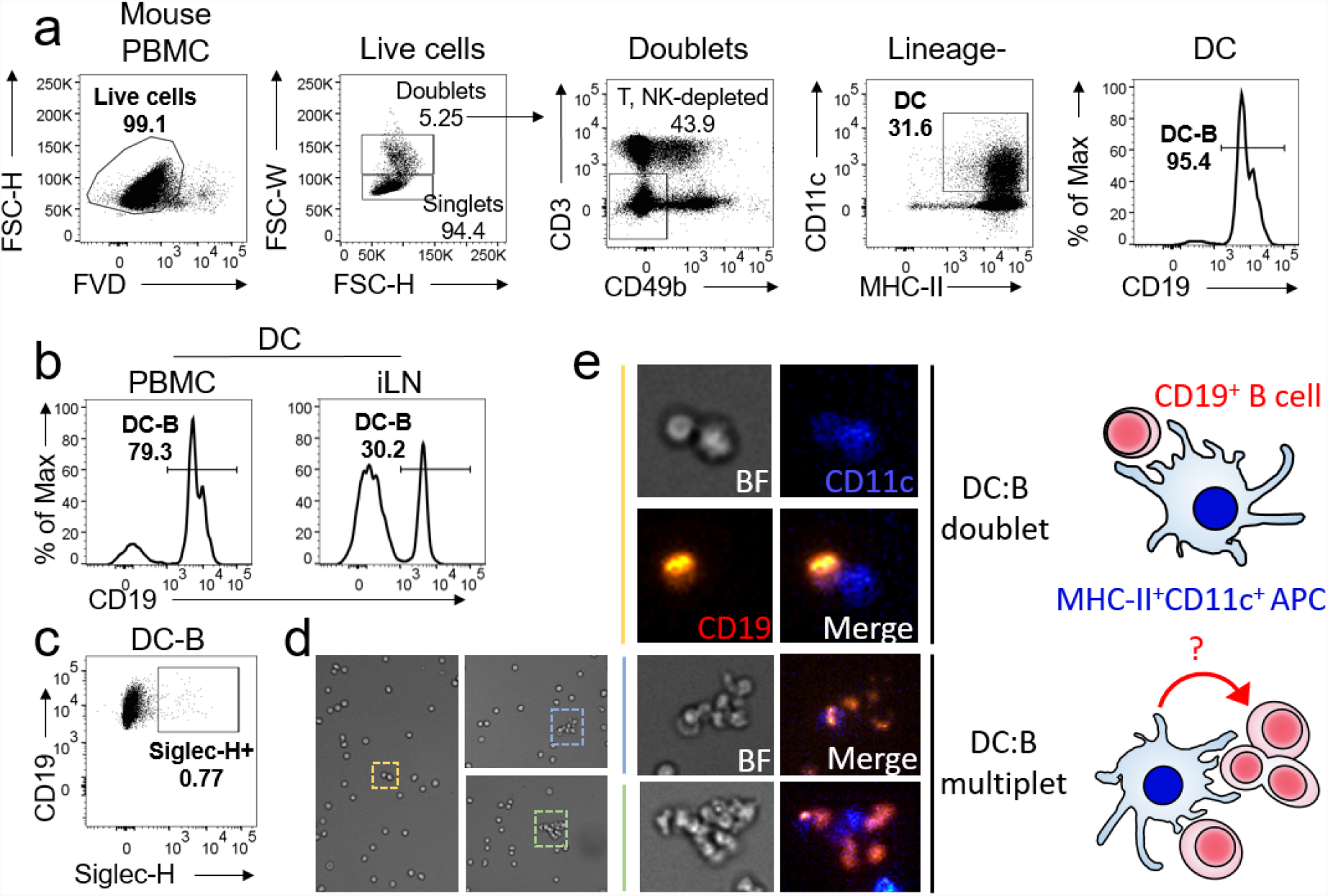
Verification of APC:B CICs in wildtype mouse blood. (a) Procedure of flow cytometry of DC:B CICs among healthy wildtype mouse blood PBMC. (b) Percentage of DC:B cell clusters among total DCs in PBMC (left) or inguinal lymph nodes (iLNs) (c) Percentage of Siglec-H+ pDCs among DC:B CICs. (d, e) microscopic images (d), close-ups images (e, left) and illustration (e, right) of DC:B doublets and multiplets.

**Figure 4.**
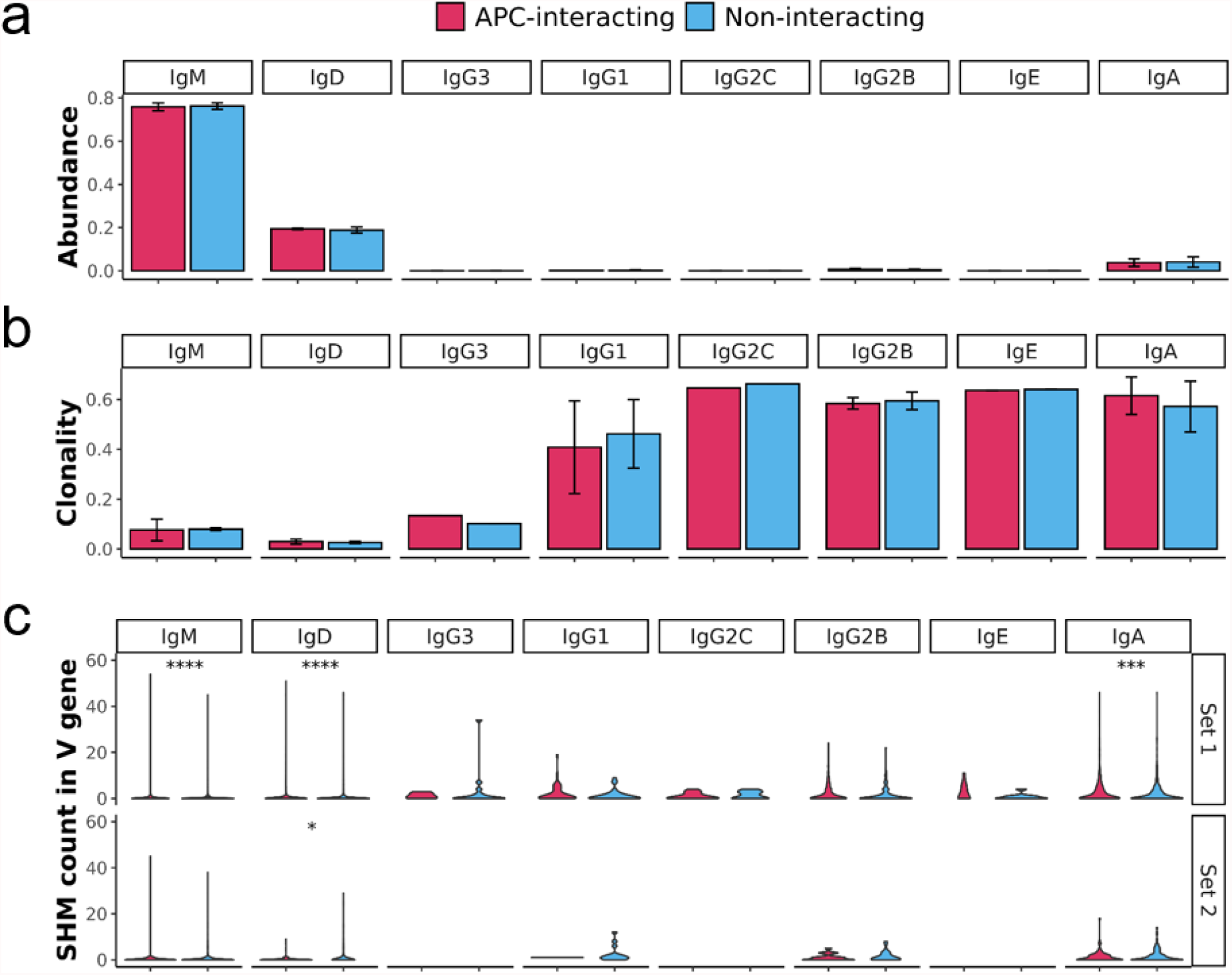
BCR profiles of APC-interacting B cells are similar to free B cells. (a) The abundance and (b) the clonality of each isotype on APC-interacting and non-interacting B cells. The mean value was plotted as a bar with the standard deviation. The clonality was measured using a Gini index, and higher clonality represents the lower diversity of BCR sequences. (c) The number of SHM in V gene of each isotype on APC-interacting and non-interacting B cells. The significance of the difference in SHM count was tested using Wilcoxon rank sum test (* : p ≤ 0.05, ** : p ≤ 0.01, *** : p ≤ 0.001, **** : p ≤ 0.0001).

Compared to DCs, APC:B CICs showed enriched CCR7 expression (**Figure S1. B**). CCR7 is a chemokine receptor that are upregulated in lymph-node migrating T cells, B cells, or DCs^24^. Whether this CCR7 expression in APC:B is from the APC or B cell or both is not clear. But it is plausible that these APC:B CICs might have higher potential for lymph-node migration compared to non-B cell-bound APCs with low or no detectable CCR7 expression, which was the majority of DCs in the PBMC datasets (**Figure S1. B**).

To test whether this APC:B CIC was a unique event limited to this dataset or not, we observed other healthy human PBMC scRNA-seq datasets from other sources^26,27^. We repeatedly detected such B cell signature-containing APCs in PBMC datasets obtained from not only 10x Genomics scRNA-seq data but also in data obtained using flow cytometry-sorted cells and microwell array-sorted cells (**Figure 2.B, C**), thus eliminating the hypothesis of APC:B CICs appearing due to single cell sequencing platform artefact. It is also noteworthy that these B cell-specific transcripts were enriched in specific APC subtypes like CD14^+^ monocytes, pDCs or CD1c+ DCs and rarely found in CD16^+^ monocytes, CLEC9A+ DCs, asDCs (**Figure 1, B, Figure 2. B, Figure S1. C**). This indicates that these APC-B cell mixed transcriptomes are not likely to have appeared by i) B cell free RNA contamination, neither ii) accidental B cell co-sorting because if so, B cell transcripts would appear evenly across APC cell types.

### Observation and BCR analysis of APC:B CICs among PBMCs in mouse

We tested whether these APC:B CICs exist in wildtype mouse PBMCs. For this, we used a sorting scheme to specifically target doublet or multiplet cells and analyze whether a subset of such multiplet contain simultaneous DC marker (CD11c, MHCII) and B cell marker (CD19) (**Figure 3.A**). Flow cytometry results showed that among cell doublet/multiplet containing DCs (CD11c+MHCII+), around 86.9% of them contained B cell (CD19+) as well (**Figure S2**). Noticeably, frequency of B cell-interacting DCs among total DCs was higher in blood than in lymph nodes (**Figure 3.B**). Also, these B cell-interacting B cells were mostly Siglec-H^-^ DCs (**Figure 3.C**).

It is known that FACS tend to produce erroneous doublet reads at low frequency. This phenomena happens when two or more separate cells are accidently encapsulated into the same droplet and imaged as a single instant and tends to increase in frequency with increased flow rate. Since we were forced to use high flow rate due to the extremely low frequency of APC:B CICs among B cell singlets, false positive cell doublets could have been the reason for our APC:B doublet observations. To determine whether this was the case or not, we imaged the flow cytometry-sorted cells using fluorescence microscopy. Definitively, cell doublets and multiplets containing CD19+ B cells and CD11c+ DCs were observable (**Figure 3. D, E**). Conclusively, we verified that DC:B CICs exist in blood.

We next asked whether the B cells under direct contact with DCs have distinct immune profile compared to non-interacting B cell population. For this, we sorted healthy mouse blood B cell singlets and DC:B CICs separately, performed RT-PCR against BCR kappa heavy chains, and performed next BCR-seq. The result showed that BCR repertoire of DC:B CICs and B cells are similar in isotype composition, clonal diversity, and V-gene somatic hyper-mutation (SHM) rates (**Figure 3**). This implied that DC-to-B cell binding is not biased towards specific BCR sequences and might occur generally across diverse B cell population.

### Dynamic changes of APC:B CICs in COVID-19 patient blood

We analyzed a unique COVID-19 patient PBMC scRNA-seq that pre-enriched APCs using MACS ^27^. Importantly, they added EDTA during APC enrichment to dissociate potential cell clusters like DC-T cell clusters. This also effectively prevents cation charge-based nonspecific cell-cell clustering. In accordance with our previous analyses, APC:B CIC-like signatures were prevalent in these APCs (**Figure 5.A, B**). Interestingly, expression level of a number of B cell subtype marker genes differed between healthy donors and COVID-19 patients (**Figure 5. B, C**). Frequencies of cells enriched with MS4A1 (CD20, expressed on naïve B cell and memory B cell) were relatively high in healthy donors than COVID-19 patients. On the contrary, frequencies of cells enriched with MZB1 or IGHG1 (markers of plasmablast or plasma cells) was higher in COVID-19 patients. MHC-II expression level decreased as disease severity increased as reported. Noticeably, APCs from moderate COVID-19 patients showed high levels of IGHM and MZB1 while severe patients showed high level of IGHG1 and lower MZB1 level. Potential interpretation of these results is as follows; blood circulating APCs, in healthy state, form clusters with CD20^+^ naïve B cells that express low levels of IgM but in pathogen (SARS-CoV-2)-infected state, they either i) stimulate the contacting naïve B cells into CD20 ^-^MZB1^+^ IgM+/IgG+ plasmablasts or ii) dissociate with naïve B cells and form stronger bonding with already active plasmablasts.

**Figure 5.**
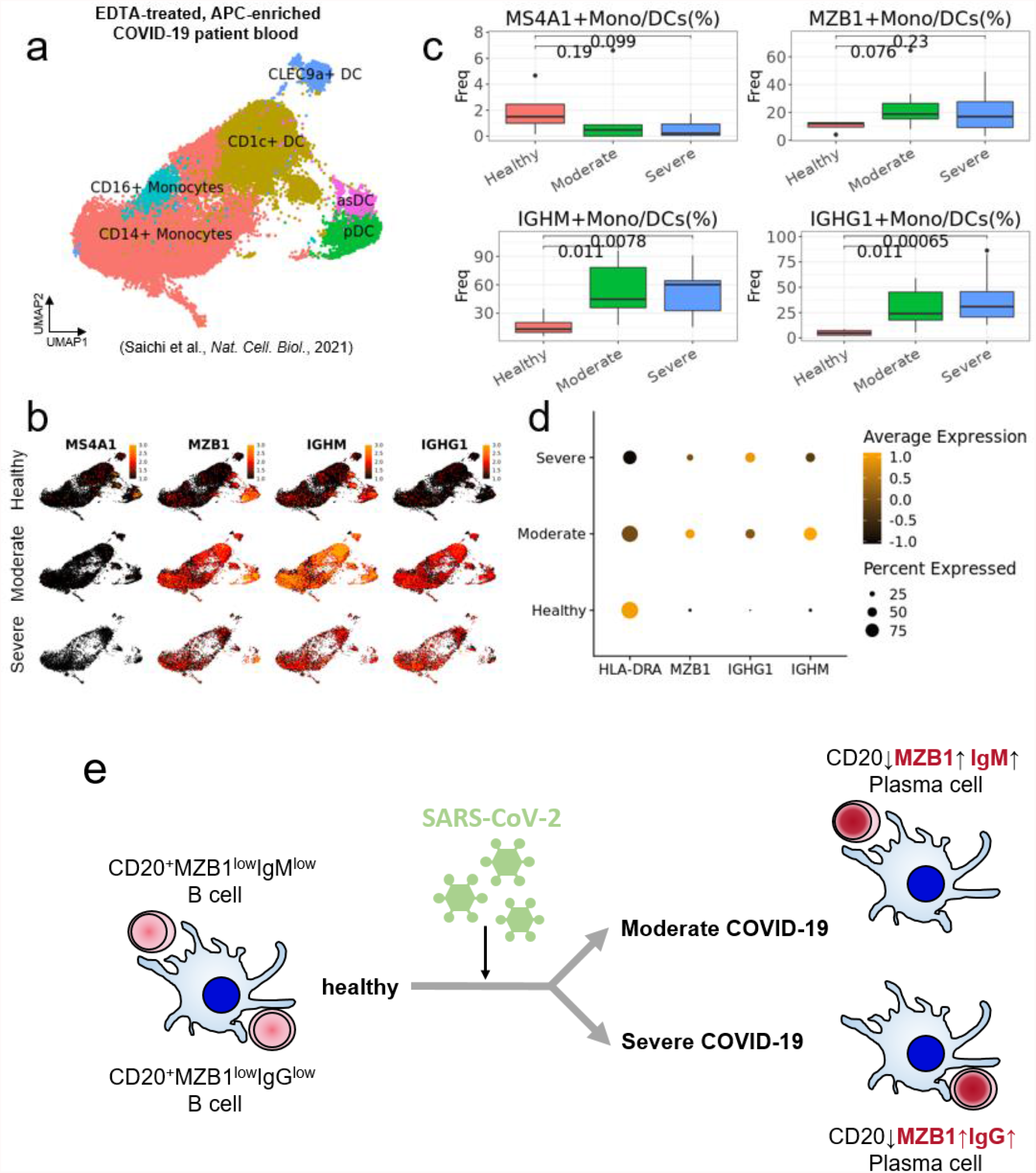
Dynamic changes of APC:B CIC-like signatures in SARS-CoV-2 infected patients. (a) UMAP plot of APC celltype clusters obtained from COVID-19 patients and healthy donors. (b) UMAP gene expression plot of selected naïve B cell and plasmablast markers among healthy donors, moderate COVID-19 patients and severe COVID-19 patients. (c) boxplots of percentage of B cell marker gene-positive APCs among total APCs. p-values were calculated using Wilcoxon rank-sum test. (d) Dotplot of B cell marker enrichment level. Dot size indicates percentage of gene-positive cells. Dot color indicates average gene expression. (e) Illustration of predicted dynamic changes of APC:B CICs in COVID-19 patients.

## Discussion

Although there were known cases of direct contact between DCs and B cells, these were known to happen in lymphoid organs. These cases involved plasmacytoid DCs (pDCs) and were shown to regulate autoreactive B cell^12^ or stimulate proliferation of B cells and immunoglobulin secretion. The latter B cell stimulation was discovered to be dependent on interaction of CD70 on DCs and CD27 on B cells, which could happen independent of type I interferon and IL-6 secretion^14^. Corroborating evidence of potential autoimmunogenic effects of pDC-B interaction spurs further question such as whether such DC contact-based B cell activation is a non-discriminatory process or biased towards certain B cell clone(s) like autoreactive B cells. We anticipate future immune profiling and antibody panning experiments with *in vitro* autoantigen-stimulated DC-B CICs will provide answers.

What subtype of B cells interact with these APC needs further investigation. Our analysis shows that CD27 -a mature B cell marker - and AIM2-a memory B cell marker-were significantly expressed in APC:B CICs. But the expression levels were not exclusive to APC:B CICs compared to other DC clusters, so the evidence is not conclusive. As mentioned above, B cells that physically interact with APCs tended to have a marginal B cell-like IgM^high^IgD+CD27+ phenotype, which implies they previously passed through germinal centers. Whether this is true could be assessed by Bcl6 mutation test^28^.

What type of immune synapse or ligand-receptor interaction (i.e. LFA-1:ICAM-1) regulates this APC:B interaction is an open question. Our BCR profiling results suggest that this interaction is not dependent to BCR sequence. Our current study was limited to healthy subjects, COVID-19 patients, or mice with no BCR repertoire comparison between healthy and disease patients. So comparing APC:B CICs (single cell transcriptome and BCR repertoire) obtained from autoimmune/infection patients and healthy subjects would be highly informative. Recent study showed that human mo-DCs are capable of antigen cross-presentation and can produce co-stimulatory signals for the induction of effector cytotoxic CD8+ T cells^21^. And it was previously shown that DCs can nondegradatively recycle endocytosed, antigen-presenting Fcγ receptors, thus enabling cell surface interaction of antigen-presenting Fcγ receptors with BCR on B cells^29^. So one could hypothesize that mo-DCs might interact with B cells via the Fcγ-antigen-BCR axis. Whether Th cells are involved in the initial formation of these APC:B clusters is uncertain at this point. Future flow cytometry examination for APC:Th:B triple-celltype cluster and putative dynamic changes of their cell-cell contact pattern (e.g. APC:Th:B to APC:B) would be needed to decipher formation mechanism of these CICs.

APCs in COVID-19 patient blood displayed B cell subtype signatures differing from healthy donors and showed correlating with disease severity. Such increased APC:IgM^high^plasmablast CIC frequency in moderate patients and increased APC:IgG^high^plasmablast CIC frequency in severe patients implies plausible immunological functions of these CICs.

Overall, we think our present study demonstrated the potential usefulness of interpreting scRNA-seq data with in mind of physically-interacting cell clusters, alongside other analysis tools, for deciphering biologically or clinically relevant information.

