## supplementary figures for "Existence of blood circulating immune-cell clusters (CICs) comprising antigen-presenting cells and B cells"

### Supplementary information

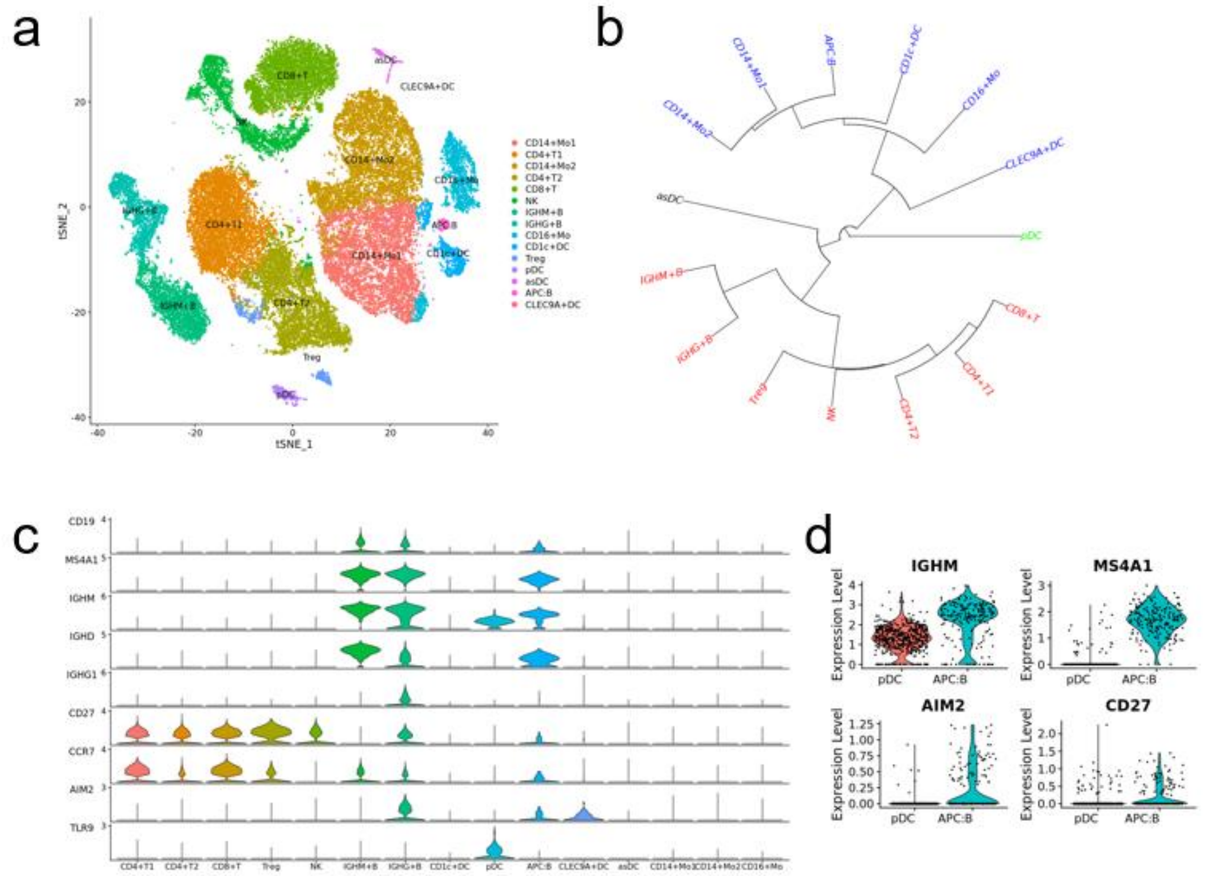


**Figure S1.** B cell-related gene expression patterns among healthy human PBMC celltypes. (a) t-SNE plot of celltype-identified clusters in healthy human PBMC 10x Genomics datasets (same data from figure 1.). (b-d) Hierarchical clustering plot (b) and violin plot of selected B-cell markers (c,d) against each celltype in (a).

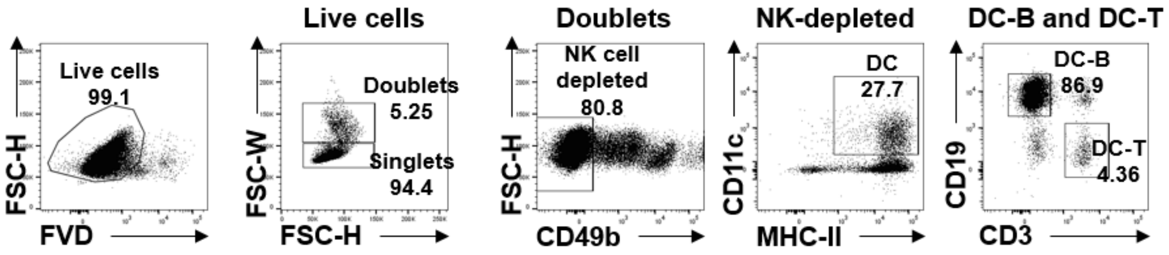


**Figure S2.** Mouse PBMC DC-B sorting strategy for measuring the percentage of DC-Bs among DCs
